## Supplementary Information for "Differential reinforcement encoding along the hippocampal long axis helps resolve the explore/exploit dilemma"

**Contents:**

**Supplemental Results.** PH/AH effects on exploration/exploitation are not explained by behavioral confounds, differences in performance, modeling choices, or effects of responses in other regions.

**Table S1.** Clusters of voxels significantly modulated by reward prediction errors (RPEs) in whole-brain analyses.
**Table S2.** Clusters of voxels significantly modulated by model-estimated entropy in whole-brain analyses.
**Table S3.** Factor structure of regions responsive to reward prediction error.
**Table S4.** Factor structure of regions responsive to entropy.
**Table S5.** Behavioral effects of hippocampal signals: main and sensitivity analyses, fMRI session.
**Table S6.** Behavioral effects of hippocampal signals: main and sensitivity analyses, replication session.

**Figure S1.** Posterior hippocampal responses to reward prediction error in the current study but not in prior meta-analyses.
**Figure S2.** Encoding of reinforcement along the A-P axis when separate regressors are used to model the first half versus second half of each run of the clock task.
**Figure S3.** Hippocampal encoding of immediately preceding response time: sensitivity test for multi-level analysis of deconvolved responses.
**Figure S4.** Encoding of reinforcement along the A-P axis when modeled by simultaneous versus separate model-based decision signals in voxelwise analyses.
**Figure S5.** Hippocampal online responses to the global value maximum, sensitivity analysis using a restrictive mask.
**Figure S6.** Hippocampal encoding of reinforcement, sensitivity analysis using a restrictive mask.

### Supplemental Results

#### PH/AH effects on exploration/exploitation are not explained by behavioral confounds, differences in performance, modeling choices, or effects of responses in other regions

In sensitivity analyses, we ascertained that the doubly dissociable effects of PH vs. AH responses on exploration vs. exploitation remained unchanged after controlling for behavioral variables (trial, contingency, maximum available value, uncertainty, and their interactions), subject-level performance (mean entropy and value) and for interactions between these potential confounds and hippocampal responses (Table S5). The addition of each block of variables improved the fit of the model predicting RT but did not change the substantive effects of AH and PH on choice. Sensitivity analyses of the replication session yielded essentially identical results such that PH predicted exploration (RT swing) and AH predicted exploitation (convergence on global value maximum) even after controlling for other behavioral variables (Table S6).

Since we used the SCEPTIC RL model to generate RPE and entropy estimates tested against hippocampal activity, we ascertained that our findings were not tautologically explained by other model-derived covariates (e.g. maximum available value) into statistical models predicting behavior. We thus performed "model-free" (i.e., no variables from the SCEPTIC model) behavioral analyses including only $RT_{t-1}$, trial, last outcome and condition as covariates. In these reduced models, PH RPE still strongly predicted greater exploration (fMRI session, $RT_{t-1}$ × PH: $t = -12.46$, $p < 10^{-15}$; replication session, $t = -7.32$, $p < 10^{-12}$), particularly following rewards (fMRI session, $RT_{t-1}$ × last outcome × PH: $t = 5.77$, $p < 10^{-8}$; replication session, $t = 4.27$, $p < 10^{-4}$). In a model without $RT_{Vmax}$, AH low entropy responses predicted less exploration (fMRI session, $RT_{t-1}$ × AH: $t = 2.92$, $p = .004$; replication session, $t = 1.30$, $p = .20$), particularly after reward omissions (fMRI session, $RT_{t-1}$ × last outcome × AH: $t = 2.18$, $p = .029$; replication session, $t = 5.15$, $p < 10^{-6}$). However, adding $RT_{Vmax}$ back to the model revealed that this effect was partly explained by greater exploitation in participants with strong AH responses (fMRI session, $RT_{t-1}$ × AH: $t = 0.81$, $p = .41$, $RT_{t-1}$ × last outcome × AH: $t = 2.19$, $p = .028$, $RT_{Vmax}$ × AH: $t = 2.74$, $p = .006$, -1/trial × $RT_{Vmax}$ × AH: $t = 2.44$, $p = 0.015$; replication session, $RT_{t-1}$ × AH: $t = 0.57$, $p = .566$, $RT_{t-1}$ × last outcome × AH: $t = 5.01$ $p < 10^{-6}$, $RT_{Vmax}$ × AH: $t = 1.48$, $p = .14$, -1/trial × $RT_{Vmax}$ × AH: $t = 3.07$, $p = 0.002$). Finally, since participants' hippocampal activity estimates were based on reinforcement, we removed all reinforcement information from the model predicting choices. In this reduced model, our primary effects of interest were still robust, even though the effect of AH on exploitation during the replication session was only evident later in learning (fMRI session, $RT_{t-1}$ × PH: $t = -9.53$, $p < 10^{-15}$, $RT_{Vmax}$ × AH: $t = 2.63$, $p = .009$, -1/trial × $RT_{Vmax}$ × AH: $t = 2.53$, $p = 0.011$; replication session, $RT_{t-1}$ × PH: $t = -3.97$, $p < 10^{-4}$, $RT_{Vmax}$ × AH: $t = 0.80$, $p = 0.42$, -1/trial × $RT_{Vmax}$ × AH: $t = 3.01$, $p = 0.003$).

Given the established role of cortico-striatal networks in reward learning, it was important to ascertain that hippocampal signals predicted behavior above and beyond cortico-striatal signals identified in the literature and our study in particular (Tables S1-S4). Thus, we tested for region and signal specificity of hippocampal effects on exploration/exploitation after controlling for the strength of neural reward signals in other key regions. These included vmPFC expected value responses, dorsal attention network responses to high entropy and cortico-striatal prediction error responses. All effects of hippocampal signals were robust in both the fMRI and replication sessions (fMRI session: $RT_{t-1}$× PH: $t = -7.16$, $p < 10^{-12}$, $RT_{Vmax}$× AH: $t = 3.57$, $p < .001$, -1/trial × $RT_{Vmax}$ × AH: $t = 2.07$, $p = 0.038$; replication session: $RT_{t-1}$× PH: $t = -5.45$, $p < 10^{-7}$, $RT_{Vmax}$× AH: $t = 3.17$, $p = .0015$, -1/trial × $RT_{Vmax}$ × AH: $t = 2.52$, $p = 0.012$).

**Table S1.** Clusters of voxels significantly modulated by reward prediction errors (RPEs) in whole-brain analyses.

| | | | | Cluster center-of-mass MNI coordinates | | |
|---|---|---|---|---|---|---|
| | **Regions within cluster** | **Number of Voxels** | $z_{max}$ | **x** | **y** | **z** |
| 1 | Dorsal anterior cingulate cortex, midcingulate cortex, bilateral fusiform gyrus, bilateral intraparietal sulcus, bilateral supplementary motor area, bilateral cerebellum (Crus I), bilateral precuneus | 13641 | 8.58 | 0.7 | -56.0 | 15.2 |
| 2 | Left ventral striatum, right ventral striatum, left putamen, right putamen, bilateral thalamus, left inferior frontal gyrus pars opercularis, left anterior insula, right anterior insula | 3803 | 8.03 | -7.4 | 0.7 | 4.9 |
| 3 | Left inferior frontal gyrus pars triangularis | 704 | 5.15 | -42.0 | 37.0 | 9.2 |
| 4 | Right inferior frontal gyrus pars opercularis | 494 | 7.58 | 47.8 | 10.2 | 26.6 |
| 5 | Right inferior frontal gyrus pars triangularis | 431 | 5.14 | 43.6 | 38.4 | 12.6 |
| 6 | Left anterior insula/frontal operculum | 204 | 5.40 | -33.2 | 21.6 | -1.6 |
| 7 | Right posterior hippocampus | 190 | 7.21 | 21.8 | -34.2 | -1.5 |
| 8 | Left superior occipital gyrus | 185 | -5.89 | -13.9 | -97.4 | 18.2 |
| 9 | Right superior occipital gyrus | 165 | -4.53 | 18.3 | -94.8 | 20.2 |
| 10 | Left posterior hippocampus | 152 | 5.24 | 20.8 | -36.1 | -0.8 |
| 11 | Right superior frontal gyrus | 146 | 4.43 | 28.8 | 4.9 | 62.0 |

*Note.* Clusters of significant voxels were identified based on a voxelwise threshold of $|z| > 3.09$ ($p < .001$, one-tailed) and a cluster threshold of $p < .05$. The cluster thresholds were identified using by randomizing the sign of the residuals of a one-sample *t*-test of the whole-brain RPE analysis using AFNI *3dttest++ -Clustsim*. This procedure does not assume a Gaussian autocorrelation function and determined that clusters > 107 voxels were significant at the whole-brain level. Cluster numbers are given in the order of decreasing size.

**Table S2.** Clusters of voxels significantly modulated by model-estimated entropy in whole-brain analyses.

| | | | | Cluster center-of-mass MNI coordinates | | |
|---|---|---|---|---|---|---|
| | **Regions within cluster** | **Number of Voxels** | $z_{max}$ | **x** | **y** | **z** |
| | *Regions sensitive to low entropy* | | | | | |
| 4 | Left postcentral gyrus | 417 | -4.53 | -38.9 | -24.1 | 61.6 |
| 5 | Bilateral ventromedial prefrontal cortex | 282 | -4.39 | -6.2 | 52.5 | -5.9 |
| 8 | Right Rolandic operculum | 193 | -4.98 | 51.2 | -20.5 | 16.7 |
| 9 | Left anterior hippocampus | 159 | -5.26 | -28.3 | -16.7 | -16.9 |
| 10 | Left fusiform gyrus | 148 | -5.02 | -22.8 | -44.9 | -13.8 |
| 11 | Left Rolandic operculum | 132 | -4.88 | -52.5 | -13.3 | 16.5 |
| | *Regions sensitive to high entropy* | | | | | |
| 1 | Bilateral precuneus, bilateral inferior parietal lobule | 2435 | 6.64 | 10.1 | -60.6 | 54.7 |
| 2 | Right superior frontal gyrus/frontal eye field | 664 | 5.70 | 26.6 | 1.9 | 56.1 |
| 3 | Left cerebellum (Crus 1) | 440 | 5.14 | -37.6 | -55.6 | -34.6 |
| 6 | Left superior frontal gyrus/frontal eye field | 263 | 5.36 | -26.0 | -2.2 | 58.1 |
| 7 | Right rostrolateral prefrontal cortex | 208 | 4.97 | 33.3 | 56.6 | 6.2 |

*Note.* Clusters of significant voxels were identified based on a voxelwise threshold of $|z| > 3.09$ ($p < .001$, one-tailed) and a cluster threshold of $p < .05$. The cluster thresholds were identified using by randomizing the sign of the residuals of a one-sample *t*-test of the whole-brain RPE analysis using AFNI *3dttest++ -Clustsim*. This procedure does not assume a Gaussian autocorrelation function and determined that clusters > 117 voxels were significant at the whole-brain level. Cluster numbers are given in the order of decreasing size.

**Table S3.** Factor structure of regions responsive to reward prediction error

| | Region | Factor 1 | Factor 2 |
|---|---|---|---|
| 4 | Right inferior frontal gyrus pars opercularis | **0.887** | 0.006 |
| 5 | Right inferior frontal gyrus pars triangularis | **0.878** | -0.151 |
| 6 | Left anterior insula/frontal operculum | **0.740** | 0.052 |
| 2 | Bilateral striatum | **0.645** | 0.294 |
| 1 | Cingulate cortex and precuneus | **0.622** | 0.169 |
| 3 | Left inferior frontal gyrus pars triangularis | **0.603** | 0.115 |
| 11 | Right superior frontal gyrus | **0.582** | -0.09 |
| 10 | Left posterior hippocampus | -0.04 | **0.973** |
| 7 | Right posterior hippocampus | 0.196 | **0.731** |

Note. The highest factor loading is printed in bold face. Cluster numbers match Table S1.

**Table S4.** Factor structure of regions responsive to entropy

| | Region | Factor 1 | Factor 2 |
|---|---|---|---|
| 10 | Left fusiform gyrus | **0.830** | 0.213 |
| 8 | Right Rolandic operculum | **0.796** | 0.304 |
| 11 | Left Rolandic operculum | **0.717** | 0.298 |
| 9 | Left anterior hippocampus | **0.710** | -0.066 |
| 4 | Left postcentral gyrus | **0.708** | 0.241 |
| 5 | Bilateral ventromedial prefrontal cortex | **0.538** | -0.199 |
| 1 | Bilateral precuneus | 0.29 | **0.850** |
| 2 | Right superior frontal gyrus | 0.142 | **0.831** |
| 3 | Left cerebellum (Crus 1) | 0.057 | **0.742** |
| 6 | Left superior frontal gyrus | 0.349 | **0.638** |
| 7 | Right rostrolateral prefrontal cortex | -0.133 | **0.632** |

*Note.* The highest factor loading is printed in bold face. Cluster numbers match Table S2.

**Table S5.** Behavioral effects of hippocampal signals: main and sensitivity analyses, fMRI session.

| | Dependent variable: | | | |
|---|---|---|---|---|
| | RT | | | |
| | Main analysis | + contingency | + choice uncertainty | + subject-level performance |
| | (1) | (2) | (3) | (4) |
| -1/trial | 0.01 (0.01) | -0.01 (0.03) | 0.02 (0.03) | 0.02 (0.03) |
| RT(t-1) | 0.5 (0.01)$^{***}$ | 0.5 (0.01)$^{***}$ | 0.5 (0.01)$^{***}$ | 0.5 (0.01)$^{***}$ |
| RT(Vmax, t-1) | 0.1 (0.01)$^{***}$ | 0.1 (0.01)$^{***}$ | 0.1 (0.01)$^{***}$ | 0.1 (0.01)$^{***}$ |
| last outcome: omission vs. reward | -0.2 (0.01)$^{***}$ | -0.2 (0.01)$^{***}$ | -0.2 (0.01)$^{***}$ | -0.2 (0.01)$^{***}$ |
| Vmax, within-subject | 0.01 (0.01) | 0.01 (0.01) | 0.01 (0.01) | 0.01 (0.01) |
| entropy, within-subject | 0.02 (0.01)$^{***}$ | 0.02 (0.01)$^{***}$ | 0.02 (0.01)$^{*}$ | 0.02 (0.01)$^{*}$ |
| AH low entropy resp. | -0.01 (0.1) | -0.1 (0.2) | -0.1 (0.2) | 1.2 (0.6) |
| PH RPE resp. | -0.03 (0.04) | -0.1 (0.04) | -0.1 (0.04) | -0.2 (0.1) |
| contingency: CEVR vs. CEV | | 0.1 (0.05) | 0.05 (0.05) | 0.1 (0.05) |
| contingency: DEV vs. CEV | | -0.1 (0.04)$^{***}$ | -0.1 (0.04)$^{***}$ | -0.1 (0.04)$^{**}$ |
| contingency: IEV vs. CEV | | 0.1 (0.04)$^{***}$ | 0.1 (0.04)$^{***}$ | 0.1 (0.04)$^{***}$ |
| uncertainty of last choice | | | 0.03 (0.01)$^{***}$ | 0.03 (0.01)$^{***}$ |
| mean entropy, between-subjects | | | | 0.6 (0.1)$^{***}$ |
| mean Vmax, between-subjects | | | | -0.001 (0.001) |
| -1/trial * RT(t-1) | -0.1 (0.01)$^{***}$ | -0.1 (0.01)$^{***}$ | -0.1 (0.01)$^{***}$ | -0.1 (0.01)$^{***}$ |
| -1/trial * last outcome | 0.1 (0.01)$^{***}$ | 0.1 (0.01)$^{***}$ | 0.1 (0.01)$^{***}$ | 0.1 (0.01)$^{***}$ |
| -1/trial * RT(Vmax) | 0.01 (0.02) | 0.02 (0.02) | 0.02 (0.02) | 0.02 (0.02) |
| -1/trial * Vmax | 0.01 (0.01)$^{*}$ | 0.02 (0.01)$^{*}$ | 0.01 (0.01)$^{*}$ | 0.02 (0.01)$^{*}$ |
| -1/trial * entropy | -0.01 (0.01) | -0.01 (0.01) | -0.01 (0.01) | -0.01 (0.01) |
| -1/trial * AH | -0.01 (0.04) | 0.1 (0.1) | 0.03 (0.1) | 0.03 (0.1) |
| -1/trial * PH | 0.01 (0.01) | 0.02 (0.02) | 0.02 (0.02) | 0.02 (0.02) |
| RT(t-1) * RT(Vmax) | -0.003 (0.01) | -0.002 (0.01) | 0.003 (0.01) | 0.004 (0.01) |
| RT(t-1) * last outcome | -0.4 (0.01)$^{***}$ | -0.4 (0.01)$^{***}$ | -0.4 (0.01)$^{***}$ | -0.4 (0.01)$^{***}$ |
| RT(t-1) * Vmax | -0.03 (0.01)$^{***}$ | -0.03 (0.01)$^{***}$ | -0.03 (0.01)$^{***}$ | -0.03 (0.01)$^{***}$ |
| RT(t-1) * entropy | -0.02 (0.01)$^{**}$ | -0.02 (0.01)$^{**}$ | -0.02 (0.01)$^{**}$ | -0.02 (0.01)$^{**}$ |
| RT(t-1) * AH | 0.04 (0.04) | 0.02 (0.04) | 0.04 (0.04) | 0.04 (0.04) |
| **RT(t-1) * PH** | **-0.1 (0.01)$^{***}$** | **-0.1 (0.01)$^{***}$** | **-0.1 (0.01)$^{***}$** | **-0.1 (0.01)$^{***}$** |
| RT(Vmax) * last outcome | 0.1 (0.01)$^{***}$ | 0.1 (0.01)$^{***}$ | 0.1 (0.01)$^{***}$ | 0.1 (0.01)$^{***}$ |
| RT(Vmax) * Vmax | 0.1 (0.01)$^{***}$ | 0.1 (0.01)$^{***}$ | 0.1 (0.01)$^{***}$ | 0.1 (0.01)$^{***}$ |
| RT(Vmax) * entropy | 0.01 (0.01) | 0.01 (0.01) | 0.01 (0.01) | 0.01 (0.01) |
| **RT(Vmax) * AH** | **0.1 (0.04)$^{***}$** | **0.1 (0.04)$^{**}$** | **0.1 (0.04)$^{**}$** | **0.1 (0.04)$^{**}$** |
| RT(Vmax) * PH | 0.01 (0.01) | 0.01 (0.01) | 0.01 (0.01) | 0.01 (0.01) |
| last outcome * Vmax | -0.02 (0.01) | -0.02 (0.01)$^{*}$ | -0.02 (0.01)$^{*}$ | -0.02 (0.01)$^{*}$ |
| last outcome * entropy | -0.03 (0.01)$^{**}$ | -0.03 (0.01)$^{**}$ | -0.03 (0.01)$^{**}$ | -0.03 (0.01)$^{**}$ |
| last outcome * AH | 0.2 (0.1)$^{**}$ | 0.2 (0.1)$^{**}$ | 0.2 (0.1)$^{**}$ | 0.2 (0.1)$^{**}$ |
| last outcome * PH | -0.02 (0.01) | -0.02 (0.01) | -0.02 (0.01) | -0.02 (0.01) |
| Vmax * entropy | 0.004 (0.005) | 0.005 (0.005) | 0.004 (0.005) | 0.005 (0.005) |
| Vmax * AH | -0.05 (0.02) | -0.04 (0.02) | -0.05 (0.02) | -0.05 (0.02) |
| Vmax * PH | -0.003 (0.005) | -0.003 (0.005) | -0.003 (0.005) | -0.003 (0.005) |
| Entropy * AH | 0.02 (0.02) | 0.02 (0.02) | 0.03 (0.03) | 0.03 (0.03) |
| Entropy * PH | -0.01 (0.005)$^{*}$ | -0.01 (0.005)$^{*}$ | -0.01 (0.01)$^{*}$ | -0.01 (0.01)$^{*}$ |
| AH * PH | 0.01 (0.1) | 0.01 (0.1) | 0.01 (0.1) | 0.04 (0.1) |
| -1/trial * CEVR | | 0.02 (0.04) | 0.01 (0.04) | 0.01 (0.04) |

| | | | | |
|---|---|---|---|---|
| -1/trial * DEV | | 0.02 (0.03) | 0.02 (0.03) | 0.02 (0.03) |
| -1/trial * IEV | | 0.03 (0.03) | 0.03 (0.03) | 0.03 (0.03) |
| AH * CEVR | | 0.03 (0.2) | 0.05 (0.2) | 0.005 (0.2) |
| AH * DEV | | -0.02 (0.2) | -0.02 (0.2) | -0.01 (0.2) |
| AH * IEV | | 0.2 (0.2) | 0.2 (0.2) | 0.2 (0.2) |
| PH * CEVR | | 0.05 (0.04) | 0.05 (0.04) | 0.05 (0.04) |
| PH * DEV | | 0.05 (0.03) | 0.05 (0.03) | 0.04 (0.03) |
| PH * IEV | | 0.04 (0.03) | 0.04 (0.03) | 0.05 (0.03) |
| AH * uncertainty | | | -0.1 (0.03) | -0.1 (0.03) |
| PH * uncertainty | | | 0.001 (0.01) | 0.001 (0.01) |
| AH * mean entropy | | | | -1.1 (0.7) |
| AH * mean Vmax | | | | -0.01 (0.01) |
| PH * mean entropy | | | | 0.03 (0.1) |
| PH * mean Vmax | | | | 0.001 (0.001) |
| RT(t-1) * last outcome * AH | 0.1 (0.05) | 0.1 (0.05) | 0.1 (0.05) | 0.1 (0.05) |
| **RT(t-1) * last outcome * PH** | **0.1 (0.01)*** | **0.1 (0.01)*** | **0.1 (0.01)*** | **0.1 (0.01)*** |
| **-1/trial * RT(Vmax) * AH** | **0.1 (0.05)** | **0.1 (0.05)** | **0.1 (0.05)** | **0.1 (0.05)** |
| -1/trial * RT(Vmax) * PH | 0.01 (0.01) | 0.01 (0.01) | 0.01 (0.01) | 0.01 (0.01) |
| -1/trial * CEVR * AH | | -0.2 (0.2) | -0.2 (0.2) | -0.2 (0.2) |
| -1/trial * DEV * AH | | -0.1 (0.1) | -0.1 (0.1) | -0.1 (0.1) |
| -1/trial * IEV * AH | | -0.1 (0.1) | -0.1 (0.1) | -0.1 (0.1) |
| -1/trial * CEVR * PH | | -0.02 (0.03) | -0.02 (0.03) | -0.02 (0.03) |
| -1/trial * DEV * PH | | -0.02 (0.03) | -0.02 (0.03) | -0.02 (0.03) |
| -1/trial * IEV * PH | | -0.01 (0.03) | -0.004 (0.03) | -0.005 (0.03) |
| Constant | 0.1 (0.03)** | 0.1 (0.05)* | 0.1 (0.05) | -0.5 (0.2)** |
| Observations | 27,253 | 27,253 | 27,253 | 27,253 |
| Akaike Inf. Crit. | 61,516.6 | 61,487.9 | 61,496.5 | 61,520.9 |

* p<0.05; ** p<0.01; *** p<0.001

Table s5 legend. Regression coefficient estimate (standard error). Effects of interest are bolded.

**Table S6.** Behavioral effects of hippocampal signals: main and sensitivity analyses, replication session.

| | *Dependent variable:* | | | |
|---|---|---|---|---|
| | RT | | | |
| | Main analysis | + contingency | + choice uncertainty | + subject-level performance |
| | (1) | (2) | (3) | (4) |
| -1/trial | 0.1 (0.01)*** | 0.02 (0.04) | 0.1 (0.04)* | 0.1 (0.04)* |
| RT(t-1) | 0.5 (0.01)*** | 0.5 (0.01)*** | 0.5 (0.01)*** | 0.5 (0.01)*** |
| RT(Vmax, t-1) | 0.1 (0.01)*** | 0.1 (0.01)*** | 0.1 (0.01)*** | 0.1 (0.01)*** |
| last outcome: omission vs. reward | -0.1 (0.01)*** | -0.1 (0.01)*** | -0.1 (0.01)*** | -0.1 (0.01)*** |
| Vmax, within-subject | 0.001 (0.01) | 0.01 (0.01) | 0.01 (0.01) | 0.01 (0.01) |
| entropy, within-subject | 0.04 (0.01)*** | 0.04 (0.01)*** | 0.03 (0.01)*** | 0.03 (0.01)*** |
| AH low entropy resp. | 0.04 (0.1) | 0.3 (0.2) | 0.3 (0.2) | 1.0 (0.9) |
| PH RPE resp. | -0.01 (0.03) | 0.03 (0.04) | 0.04 (0.04) | 0.1 (0.2) |
| contingency: CEVR vs. CEV | | -0.05 (0.05) | -0.05 (0.05) | -0.1 (0.05) |
| contingency: DEV vs. CEV | | -0.2 (0.04)*** | -0.2 (0.04)*** | -0.2 (0.04)*** |
| contingency: IEV vs. CEV | | 0.1 (0.04)** | 0.1 (0.04)** | 0.1 (0.04)** |
| uncertainty of last choice | | | 0.04 (0.01)*** | 0.04 (0.01)*** |
| mean entropy, between-subjects | | | | 0.3 (0.2) |
| mean Vmax, between-subjects | | | | 0.005 (0.001)** |
| -1/trial * RT(t-1) | -0.003 (0.01) | -0.01 (0.01) | -0.004 (0.01) | -0.004 (0.01) |
| -1/trial * last outcome | 0.05 (0.01)*** | 0.04 (0.01)*** | 0.04 (0.01)*** | 0.04 (0.01)*** |
| -1/trial * RT(Vmax) | 0.02 (0.02) | 0.01 (0.02) | 0.01 (0.02) | 0.01 (0.02) |
| -1/trial * Vmax | -0.003 (0.01) | -0.01 (0.01)** | -0.01 (0.01)** | -0.02 (0.01)** |
| -1/trial * entropy | -0.003 (0.01) | -0.003 (0.01) | -0.003 (0.01) | -0.003 (0.01) |
| -1/trial * AH | 0.03 (0.04) | 0.02 (0.2) | 0.04 (0.2) | 0.04 (0.2) |
| -1/trial * PH | -0.01 (0.01) | 0.003 (0.04) | -0.004 (0.04) | -0.004 (0.04) |
| RT(t-1) * RT(Vmax) | -0.02 (0.005)*** | -0.02 (0.005)*** | -0.01 (0.005)** | -0.02 (0.005)*** |
| RT(t-1) * last outcome | -0.4 (0.01)*** | -0.4 (0.01)*** | -0.4 (0.01)*** | -0.4 (0.01)*** |
| RT(t-1) * Vmax | -0.02 (0.01)** | -0.02 (0.01)** | -0.02 (0.01)** | -0.02 (0.01)** |
| RT(t-1) * entropy | 0.04 (0.01)*** | 0.04 (0.01)*** | 0.04 (0.01)*** | 0.04 (0.01)*** |
| RT(t-1) * AH | 0.02 (0.04) | 0.01 (0.04) | 0.02 (0.04) | 0.02 (0.04) |
| **RT(t-1) * PH** | **-0.04 (0.01)*** ** | **-0.04 (0.01)*** ** | **-0.04 (0.01)*** ** | **-0.04 (0.01)*** ** |
| RT(Vmax) * last outcome | 0.1 (0.01)*** | 0.1 (0.01)*** | 0.1 (0.01)*** | 0.1 (0.01)*** |
| RT(Vmax) * Vmax | 0.1 (0.01)*** | 0.1 (0.01)*** | 0.1 (0.01)*** | 0.1 (0.01)*** |
| RT(Vmax) * entropy | -0.02 (0.01)*** | -0.02 (0.01)*** | -0.02 (0.01)*** | -0.02 (0.01)*** |
| **RT(Vmax) * AH** | **0.1 (0.03)* ** | **0.1 (0.03)* ** | **0.1 (0.03)†** | **0.1 (0.03)* ** |
| RT(Vmax) * PH | 0.01 (0.01) | 0.003 (0.01) | 0.003 (0.01) | 0.003 (0.01) |
| last outcome * Vmax | -0.01 (0.01) | -0.01 (0.01) | -0.01 (0.01) | -0.01 (0.01) |
| last outcome * entropy | -0.04 (0.01)*** | -0.04 (0.01)*** | -0.04 (0.01)*** | -0.04 (0.01)*** |
| last outcome * AH | 0.1 (0.05) | 0.1 (0.05) | 0.1 (0.05) | 0.1 (0.05) |
| last outcome * PH | 0.02 (0.01)* | 0.02 (0.01)* | 0.02 (0.01)* | 0.02 (0.01)* |
| Vmax * entropy | 0.001 (0.004) | 0.002 (0.004) | 0.002 (0.004) | 0.002 (0.004) |
| Vmax * AH | -0.02 (0.02) | -0.02 (0.02) | -0.02 (0.02) | -0.02 (0.02) |
| Vmax * PH | -0.01 (0.004) | -0.01 (0.004) | -0.01 (0.004) | -0.01 (0.004) |
| Entropy * AH | 0.02 (0.02) | 0.02 (0.02) | 0.02 (0.02) | 0.02 (0.02) |
| Entropy * PH | -0.001 (0.004) | -0.000 (0.004) | 0.002 (0.005) | 0.002 (0.005) |
| AH * PH | -0.004 (0.1) | -0.01 (0.1) | -0.01 (0.1) | -0.01 (0.1) |
| -1/trial * CEVR | | 0.1 (0.04) | 0.04 (0.05) | 0.04 (0.05) |

| | | | | |
|---|---|---|---|---|
| -1/trial * DEV | | -0.01 (0.04) | -0.04 (0.04) | -0.04 (0.04) |
| -1/trial * IEV | | 0.1 (0.04)$^{*}$ | 0.1 (0.04) | 0.1 (0.04) |
| AH * CEVR | | -0.3 (0.2) | -0.3 (0.2) | -0.3 (0.2) |
| AH * DEV | | -0.4 (0.2)$^{*}$ | -0.4 (0.2)$^{*}$ | -0.4 (0.2)$^{*}$ |
| AH * IEV | | -0.3 (0.2) | -0.3 (0.2) | -0.3 (0.2) |
| PH * CEVR | | -0.1 (0.04) | -0.1 (0.04) | -0.1 (0.04) |
| PH * DEV | | -0.1 (0.03)$^{*}$ | -0.1 (0.03)$^{*}$ | -0.1 (0.04)$^{*}$ |
| PH * IEV | | -0.03 (0.03) | -0.03 (0.03) | -0.02 (0.04) |
| AH * uncertainty | | | -0.01 (0.03) | -0.01 (0.03) |
| PH * uncertainty | | | -0.01 (0.01) | -0.01 (0.01) |
| AH * mean entropy | | | | -0.7 (0.9) |
| AH * mean Vmax | | | | -0.003 (0.01) |
| PH * mean entropy | | | | -0.1 (0.2) |
| PH * mean Vmax | | | | 0.001 (0.001) |
| RT(t-1) * last outcome * AH | 0.2 (0.04)$^{***}$ | 0.2 (0.04)$^{***}$ | 0.2 (0.04)$^{***}$ | 0.2 (0.04)$^{***}$ |
| RT(t-1) * last outcome * PH | 0.03 (0.01)$^{***}$ | 0.03 (0.01)$^{***}$ | 0.03 (0.01)$^{***}$ | 0.03 (0.01)$^{***}$ |
| **-1/trial * RT(Vmax) * AH** | **0.1 (0.04)**$^{**}$ | **0.1 (0.04)**$^{**}$ | **0.1 (0.04)**$^{**}$ | **0.1 (0.04)**$^{**}$ |
| -1/trial * RT(Vmax) * PH | -0.02 (0.01)$^{**}$ | -0.02 (0.01)$^{*}$ | -0.02 (0.01)$^{*}$ | -0.02 (0.01)$^{*}$ |
| -1/trial * CEVR * AH | | 0.02 (0.2) | -0.001 (0.2) | 0.003 (0.2) |
| -1/trial * DEV * AH | | 0.03 (0.2) | 0.004 (0.2) | 0.01 (0.2) |
| -1/trial * IEV * AH | | -0.02 (0.2) | -0.1 (0.2) | -0.05 (0.2) |
| -1/trial * CEVR * PH | | 0.001 (0.04) | 0.002 (0.04) | 0.002 (0.04) |
| -1/trial * DEV * PH | | -0.03 (0.04) | -0.02 (0.04) | -0.03 (0.04) |
| -1/trial * IEV * PH | | -0.004 (0.04) | -0.004 (0.04) | -0.004 (0.04) |
| Constant | 0.1 (0.03)$^{**}$ | 0.1 (0.05)$^{**}$ | 0.1 (0.05)$^{*}$ | -0.3 (0.2) |
| Observations | 31,999 | 31,999 | 31,999 | 31,999 |
| Akaike Inf. Crit. | 68,325.0 | 68,240.9 | 68,233.9 | 68,265.0 |

* p<0.05; ** p<0.01; *** p<0.001
† p=.054

Table s6 legend. Regression coefficient estimate (standard error). Effects of interest are bolded.

**Table s7.** Response hazard as a function of time-varying value and uncertainty and their interactions with hippocampal signals, mixed-effects Cox model

| term | Estimate | Standard error | z statistic | *p* value |
|---|---|---|---|---|
| $RT_{t-1}$ | -0.491 | 0.008 | -62.521 | $< 10^{-15}$ |
| PH | 0.071 | 0.072 | 0.991 | 0.322 |
| AH | -0.087 | 0.087 | -0.996 | 0.319 |
| Trial | -0.288 | 0.031 | -9.299 | $< 10^{-15}$ |
| Contingency: CEVR vs. CEV | -0.067 | 0.024 | -2.752 | 0.006 |
| Contingency: DEV vs. CEV | 0.092 | 0.020 | 4.564 | 0.00001 |
| Contingency: IEV vs. CEV | -0.163 | 0.020 | -8.101 | $< 10^{-15}$ |
| Uncertainty | -0.339 | 0.011 | -30.901 | $< 10^{-15}$ |
| Value | 0.292 | 0.005 | 58.069 | $< 10^{-15}$ |
| $RT_{t\text{-}}1$ * PH | 0.043 | 0.007 | 6.116 | $< 10^{-9}$ |
| $RT_{t-1}$ * AH | -0.021 | 0.007 | -2.929 | 0.003 |
| Trial * CEVR | -0.005 | 0.034 | -0.160 | 0.873 |
| Trial * DEV | -0.032 | 0.028 | -1.160 | 0.246 |
| Trial * IEV | -0.090 | 0.028 | -3.244 | 0.001 |
| Trial * Uncertainty | -0.026 | 0.018 | -1.411 | 0.158 |
| Trial * Value | 0.042 | 0.008 | 4.966 | 0.000001 |
| PH * Value | -0.017 | 0.004 | -3.952 | 0.0001 |
| AH * Value | 0.037 | 0.005 | 8.138 | $< 10^{-15}$ |
| AH * Uncertainty | -0.017 | 0.007 | -2.604 | 0.009 |
| PH * Uncertainty | -0.007 | 0.007 | -1.109 | 0.268 |

Table s7 Legend. Events, $n = 27435$.

**Figure S1.** Posterior hippocampal responses to reward prediction error: detected in the present study, but not in prior meta-analyses.

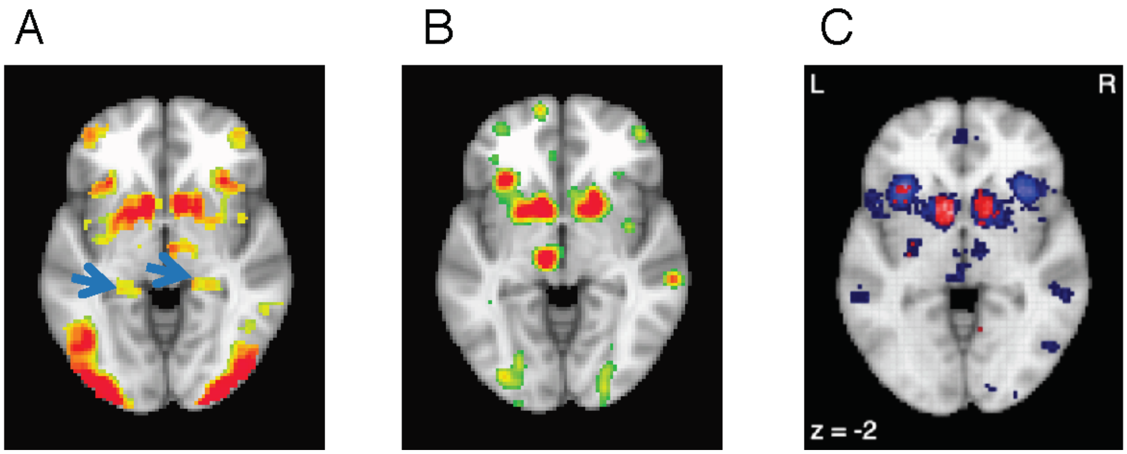


Figure S1 caption.

(A) Current study, whole-brain analysis of responses to reward prediction error, $p_{FWE\_corrected} < .05$. Blue arrows: posterior hippocampal clusters.

(B). Meta-analysis of studies using reinforcement learning models (Chase et al., 2015). Map liberally thresholded at $p_{uncorrected}<.005$.

(C) Neurosynth, “prediction error”, $p_{FDR\_corrected} < .01$.

**Figure S2.** Encoding of reinforcement along the A-P axis when separate regressors are used to model the first half versus second half of each run of the clock task.

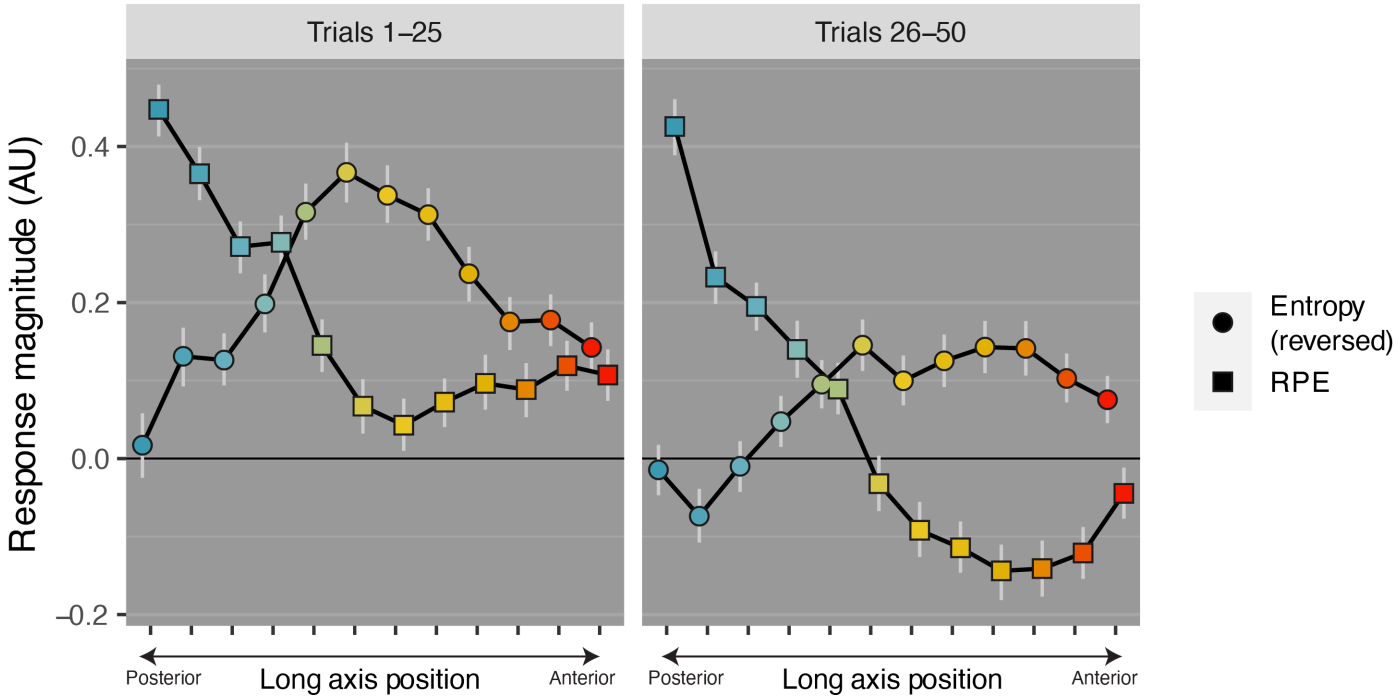


Figure S2 Caption.

Double dissociation of signals along the A-P axis when separate model-based regressors were used for the first half (Trials 1-25) versus second half (Trials 26-50) of each run of the clock task. Points represent the mean regression coefficient from voxelwise GLMs along each slice of the hippocampal long axis. The light gray vertical lines denote the bootstrapped 95% confidence interval around the mean.

**Figure S3.** Hippocampal encoding of immediately preceding response time: sensitivity test for multi-level analysis of deconvolved responses.

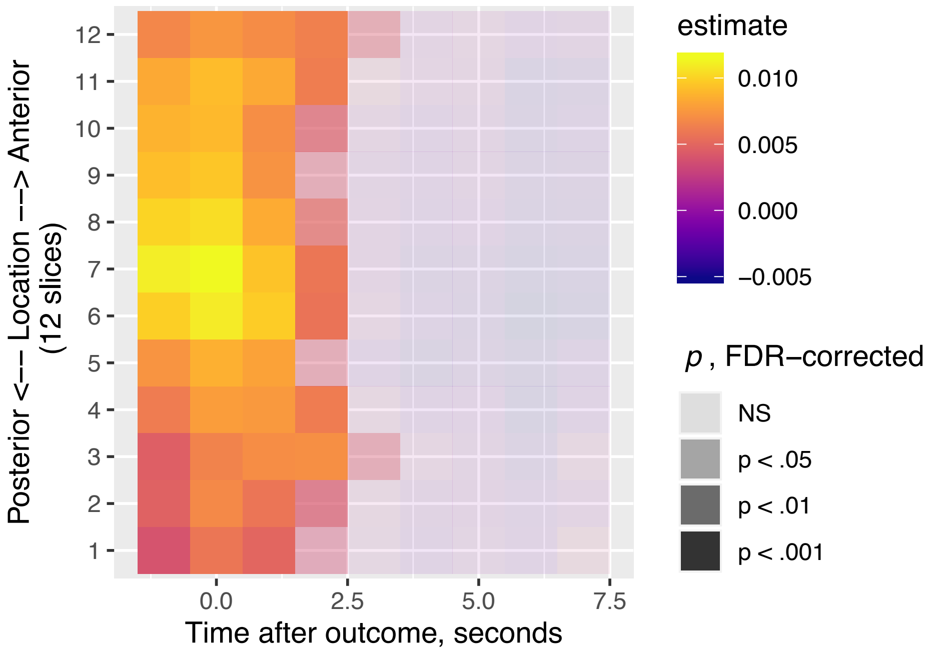


Figure S3 caption.

Unfolding hippocampal responses to the preceding response time (before current reinforcement) time-locked to current reinforcement, multi-level general linear model. Positive regression coefficients indicate stronger responses after travelling through a longer part of the interval.

**Figure S4.** Encoding of reinforcement along the A-P axis when modeled by simultaneous versus separate model-based decision signals in voxelwise analyses.

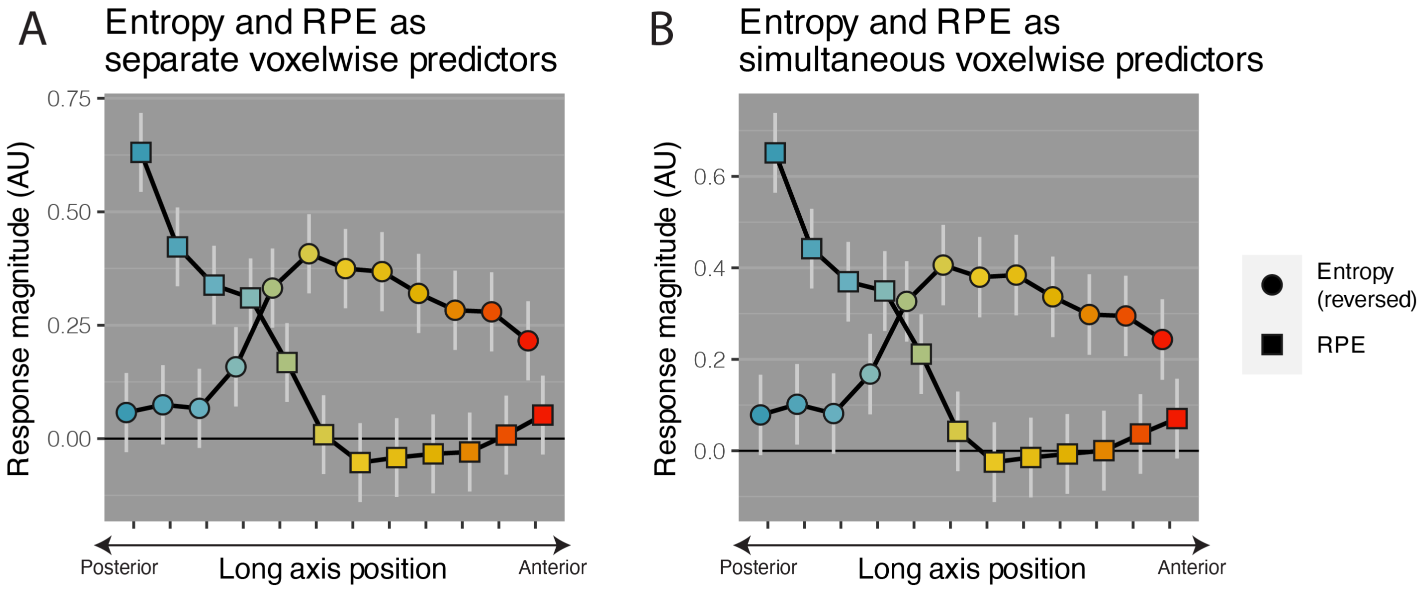


Figure S4 caption.

(A) Double dissociation of signals along the A-P axis when Entropy and RPE are entered as model-based fMRI regressors in separate GLMs. Each GLM included the clock and feedback phase regressors. Points denote the average task-related modulation in each hippocampal slice based on voxelwise regression coefficients extracted from GLM analyses. The light gray vertical lines denote the standard error from of the estimated mean from a multilevel regression model.

(B): Double dissociation of signals along the A-P axis when Entropy and RPE are entered as model-based fMRI regressors in a single GLM. The GLM also included the clock and feedback phase regressors (i.e., 4 regressors total)

**Figure S5.** Hippocampal online responses to the global value maximum, sensitivity analysis using a restrictive mask developed by Winterburn and colleagues (2013).

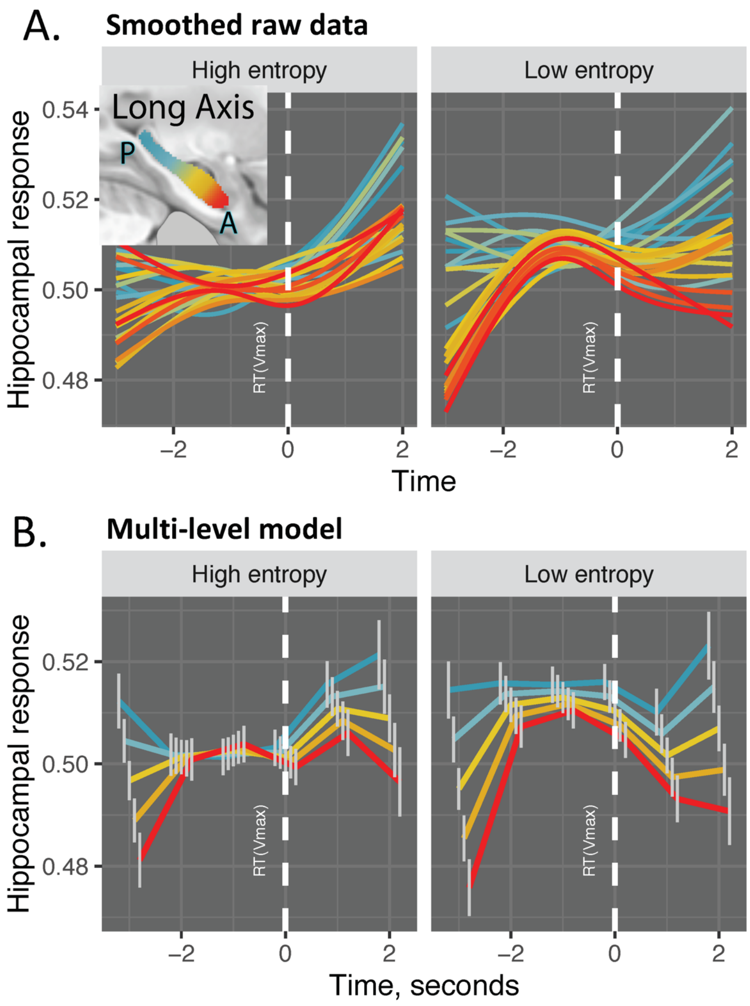


Figure S5 caption.

Analyses of deconvolved data. White dashed line: $RT_{Vmax}$

(A) Raw data, GAM smoothing with 3 knots, voxel-wise responses shown in 24 long-axis bins

(B) Multilevel model, completely general time effect. Event time × location: t = 6.95, $p < 10^{-6}$ , event time × entropy: t = 8.33, $p < 10^{-7}$. NB: since voxel-wise timeseries are normalized, only differences in the shape, but not intercept, of response can be interpreted.

**Figure S6.** Hippocampal encoding of reinforcement, sensitivity analysis using a restrictive mask.

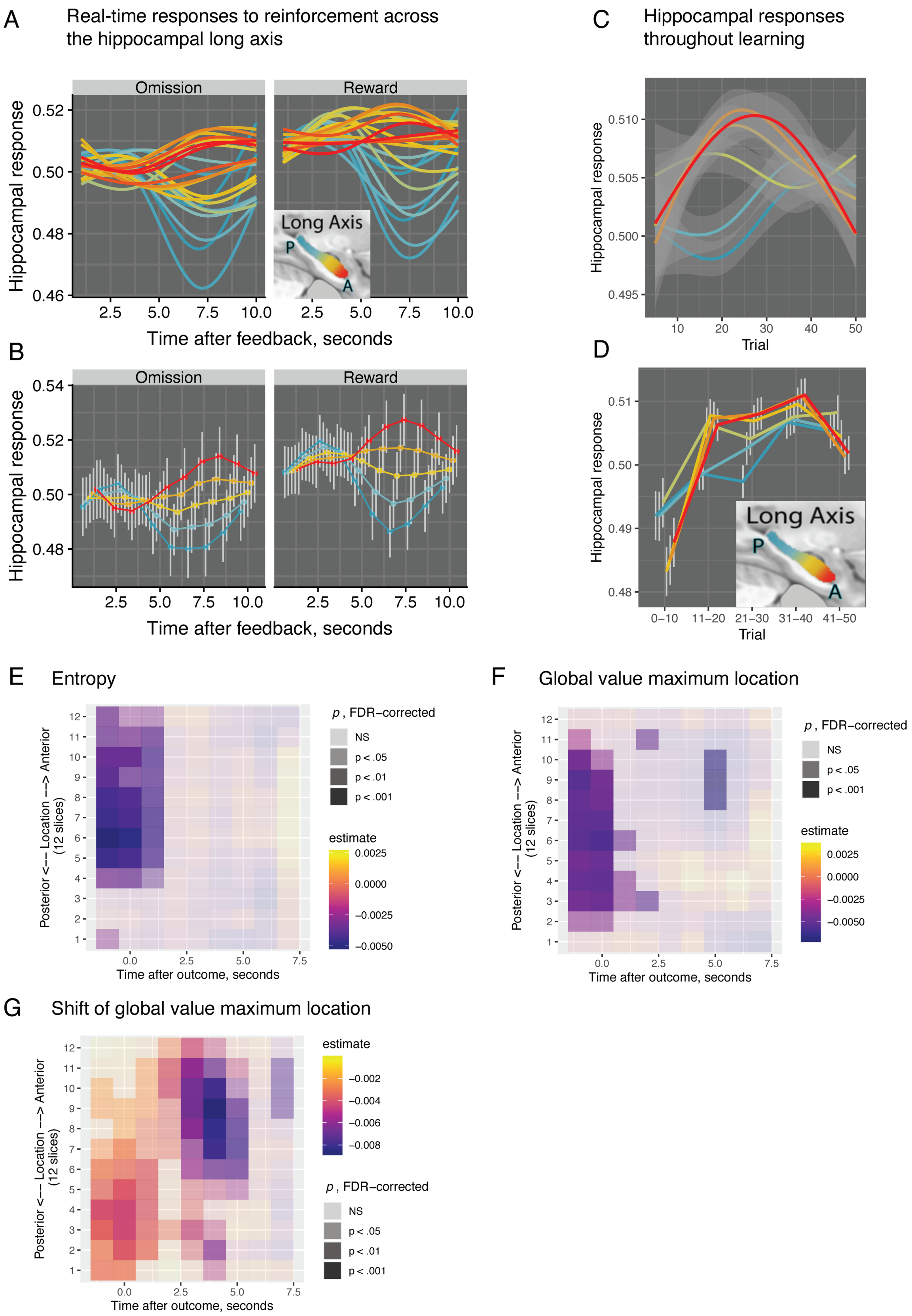


Figure S6 caption.

(A) Responses during the ITI time-locked to feedback, raw data with GAM smoothing, 3 knots.

(B) Responses during the ITI time-locked to feedback, multi-level general linear model with completely general time and bin location (12 bins) as predictors.

(C) Evolution of hippocampal responses to reinforcement across trials, raw data with GAM smoothing, 3 knots.

(D) Evolution of hippocampal responses to reinforcement across trials, multi-level general linear model with completely general *learning epoch* (five 10-trial bins) and bin location (12 bins) as predictors.

(E) Unfolding hippocampal responses to prior entropy (before current reinforcement) time-locked to current reinforcement, multi-level general linear model. Negative regression coefficients indicate stronger responses to low entropy (prominent global value maximum).

(F) Unfolding hippocampal responses to prior global value maximum location (before current reinforcement) time-locked to current reinforcement, multi-level general linear model. Negative regression coefficients indicate stronger responses to a more proximal (earlier) global value maximum.

(G) Unfolding hippocampal responses to prior the shift in the global value maximum location following current reinforcement time-locked to current reinforcement, multi-level general linear model. Negative regression coefficients indicate stronger responses when the global value maximum moves closer (earlier in the interval).
